# Pharmacological Reversal of Attention Deficits in Non-Human Primates: Implications for Alzheimer’s Disease

**DOI:** 10.1101/2025.06.02.657483

**Authors:** Casey R. Vanderlip, Shelby R. Dunn, Joseph S. Cefalu, Theresa M. Ballard, Joseph G. Wettstein, Courtney Glavis-Bloom

**Author notes:** Current affiliations ReCode Therapeutics, Menlo Park, CA 94025. Current affiliations Galwyn (UK) Ltd, Dartmouth, UK. Current affiliations TriPhase LLC, Territet, Switzerland.

## Abstract

Attention deficits emerge early in Alzheimer’s disease (AD), where cholinergic dysfunction compromises goal-directed behavior and cognitive control. Therefore, attentional impairments may serve as early indicators of cognitive decline, and also as meaningful targets for therapeutic intervention. Despite their clinical importance, attention deficits remain under-targeted by current treatments, which offer only modest benefit. To support development of more effective therapies, preclinical models that closely mirror human neurobiology and behavior are essential. Non-human primates (NHPs), with their high degree of cortical and functional similarity to humans, particularly in prefrontal regions, offer a uniquely translational platform for evaluating cognitive enhancers. We assessed pharmacological interventions targeting sustained attention using the Continuous Performance Test (CPT) in adult male cynomolgus macaques. Monkeys were trained to detect target stimuli while ignoring distractors, achieving individualized stable performance. To simulate cholinergic dysfunction, we administered scopolamine, a muscarinic acetylcholine receptor antagonist, which produced dose-dependent declines in accuracy and reaction time. Mild and severe impairment levels were identified within each animal. We then tested three compounds: nicotine, guanfacine, and donepezil. Nicotine, a nicotinic receptor agonist, fully restored performance across both impairment levels, suggesting potential benefit in both early and advanced AD. Guanfacine, an α2A adrenergic agonist, improved accuracy only under mild impairment, while donepezil, an acetylcholinesterase inhibitor, showed inconsistent effects. None of the compounds reversed scopolamine-induced slowing of reaction time, indicating specificity for attentional control. These findings highlight the utility of the NHP CPT as a pharmacologically sensitive model for detecting attentional dysfunction and evaluating pro-cognitive therapeutics in aging and neurodegeneration.

## INTRODUCTION

Attentional deficits are highly prevalent in healthy aging and become more pronounced in neurodegenerative disorders such as Alzheimer’s disease (AD) [1–3]. Although memory impairment has traditionally been the focus of AD research, attentional dysfunction emerges early in the disease course and meaningfully contributes to broader cognitive decline[3–5].

These deficits interfere with daily functioning, diminish quality of life, and increase the risk of loss of independence. Despite their clinical relevance, the mechanisms underlying attentional decline in aging and AD remain insufficiently understood.

Attention comprises several cognitive domains, including selective, divided, and sustained attention. Among these, sustained attention, or the capacity to maintain focus over time, is particularly vulnerable to age- and disease-related decline. Performance on sustained attention tasks, such as the Continuous Performance Test (CPT), declines with age and worsens further in AD, with well-documented reductions in both accuracy and response speed [6,7]. Identifying the neural and neurochemical mechanisms that support sustained attention is therefore critical for developing early interventions.

Converging evidence from neuroimaging, lesion, and electrophysiological studies implicates the prefrontal cortex (PFC) as a critical hub for sustained attention, mediating top-down control over sensory and motor systems to maintain task engagement [8–12]. Functional neuroimaging studies consistently demonstrate that sustained attention tasks activate lateral and medial PFC regions across sensory modalities [13,14]. However, the PFC is highly susceptible to aging and AD pathology, with structural and functional deterioration contributing to attentional impairments [15,16]. Reductions in PFC volume, synaptic density, and neurotransmitter availability diminish attentional capacity, while AD-related neurodegeneration further exacerbates these deficits [17,18].

Within the PFC, sustained attention critically depends on neuromodulatory inputs, particularly from the cholinergic and noradrenergic systems, both of which are highly vulnerable to aging and AD. The basal forebrain cholinergic system (BFCS), which includes the nucleus basalis of Meynert (NBM), medial septum, and diagonal band of Broca, provides the primary cholinergic innervation of the PFC [19]. This system is essential for attentional control; depletion of cholinergic signaling in rodents and non-human primates (NHPs) leads to pronounced attentional impairments [20,21]. Pharmacological blockade of cholinergic receptors (e.g., via scopolamine) impairs sustained attention [22], and in vivo microdialysis studies show increased acetylcholine (ACh) release in the PFC during sustained attention tasks [23,24]. The BFCS undergoes marked degeneration with age and in AD, with early and selective loss of NBM neurons being a hallmark of AD pathology [18,25]. Consequently, cholinergic dysfunction has been a primary target for AD therapeutics, including cholinesterase inhibitors such as donepezil [26,20].

The noradrenergic system, originating from the locus coeruleus (LC), plays a complementary role in sustaining attention, especially under conditions demanding vigilance and response inhibition [27,28]. The LC provides the brain’s sole source of cortical norepinephrine (NE), and NE depletion impairs attention across species [29,30]. Activation of post-synaptic α2A-adrenergic receptors in the PFC both through endogenous NE or pharmacological agents like guanfacine, enhances neuronal firing and improves attentional performance [31–33]. The LC is also among the earliest sites of neurodegeneration in aging and AD, with loss of noradrenergic neurons contributing to attentional and cognitive decline [34].

Importantly, aged NHPs with attentional deficits exhibit performance improvements following guanfacine administration, supporting noradrenergic modulation as a therapeutic avenue [35].

While rodent models have provided foundational insights into sustained attention, species differences in PFC structure limit their translational utility. Rodents lack a well-defined dorsolateral PFC, a region critical for executive function and attentional control in primates [36,37]. Furthermore, the architecture of neuromodulatory inputs to the PFC differs substantially between rodents and primates, complicating interpretation of preclinical findings and their translatability to humans [38].

In contrast, NHPs possess a PFC with anatomical and functional homology to humans, including similar cortical layering, connectivity, and neurotransmitter receptor distribution enabling more precise investigations into attentional mechanisms [39,40]. NHPs also exhibit complex visual attention behaviors comparable to humans, making them an ideal model for investigating sustained attention and evaluating pro-attentional therapies.

The CPT is a validated assay of sustained attention that is sensitive to aging and AD-related deficits [41,7,42]. By requiring detection of infrequent, unpredictable targets over time, the CPT minimizes working memory demands and isolates attentional function [43]. Critically, its applicability across species supports its use in translational research bridging preclinical models and clinical outcomes [44,45].

Given the critical role of sustained attention for broader cognitive function, its vulnerability to aging and neurodegeneration, and the importance of the cholinergic and noradrenergic systems in modulating sustained attention, the present study aimed to develop a CPT-based NHP model for evaluating pharmacological interventions targeting attentional deficits. We used scopolamine to model cholinergic dysfunction and tested whether pro-attentional agents including nicotine, guanfacine, and donepezil, could restore CPT performance in NHPs. By establishing a robust and translational NHP model of sustained attention, this study provides a critical platform for advancing therapeutic development in aging and AD.

## MATERIALS AND METHODS

### Subjects

Nine adult male cynomolgus macaques (*Macaca fascicularis*), approximately 6-8 years old at the onset of the study, and weighing from 6-10 kg, were used as subjects. The monkeys were singly housed in a same-sex colony room with a 12 hour light/dark cycle and at 21 ± 2 **°** C, and 40 ± 10% humidity. The subjects had access to water *ad libitum* in their home cage and were given their full daily ration of chow (Purina High Protein #5405) after daily behavioral testing was complete. All animals were also provided with daily enrichment of fresh fruit, vegetables, etc. All procedures were performed in accordance with the National Institutes of Health Guide for the Care and Use of Laboratory Animals and were approved by the SRI International Institutional Animal Care and Use Committee.

### Apparatus

Monkeys were placed into a standard Plexiglas restraint chair and then brought into a sound-attenuated chamber (30”W, 30”D, 59”H; Med Associates, St. Albans, VT) where testing took place. The chamber was outfitted with a house light, an exhaust fan for ventilation, and a white-noise generator. A panel mounted on the wall of the chamber contained a touch-sensitive monitor (Lafayette Instrument Company, Lafayette, IN) located directly in front of the monkey at a distance of approximately 30 cm, a speaker and indicator light in the upper right corner for auditory and visual reinforcement, respectively, and a hopper in the lower right corner of the panel for the monkey to receive food rewards (190 mg banana-flavored pellet; TestDiet, St. Louis, MO). Stimuli were displayed using Cambridge Neuropsychological Test Automated Battery (CANTAB®) software (Cambridge Cognition, Cambridge, MA).

### Continuous Performance Test (CPT) Procedure

The monkeys were trained 2-4 days per week on a version of the CPT [46,35] where they were ultimately required to continuously attend to a stream of 200 rectangular stimuli presented 2 seconds apart in the center of a touch screen. For training, monkeys were presented with 200 yellow rectangles (“targets”), one at a time, for 2 seconds each, and were rewarded with a banana-flavored food pellet and a “correct” auditory tone (1000 Hz, 1 sec duration) for each touch to a target (i.e., hit). Once the monkeys consistently responded to targets (hits ≥ 80%), a stepwise decrease in the percentage of target stimuli and an increase in distractor stimuli began, first by presenting 80% of the stimuli as targets and the remaining 20% as distractor until monkeys consistently touched the targets (hits ≥ 80%), and withheld responses to distractors (correct rejections ≥ 80%). The monkeys continued to receive food rewards for hits, and were punished with a 15 second time out and an “incorrect” auditory tone (40 Hz, 2 sec duration) for touching a distractor. The stepwise decrease in the percentage of stimuli that were targets continued by decreasing the instances of targets to 65%, then 50%, and finally, 30% of the total number of stimuli. Each stepwise decrease occurred when the monkeys obtained the performance criterion of ≥ 80% hits and ≥ 80% correct rejections. The final CPT on which all pharmacological testing was performed had 200 stimuli, 30% (60) of which were targets and 70% (140; 70 blue and 70 white) of which were distractors. Monkeys were considered maximally trained on the CPT when their performance maintained ≥ 80% hits and correct rejections across several testing sessions.

The data recorded were the percent of responses that were hits (correct response to a target), misses (incorrect withholding of a response to a target), correct rejections (correct withholding of a response to a distractor), false alarms (incorrect response to a distractor), and reaction time for hits (in milliseconds).

### Drug Administration

Drug doses and pre-treatment (PT) times were selected based on available literature and internal pharmacokinetic data, and attempted to target pharmacologically relevant doses in relation to human clinical studies. Donepezil (0.3-3.0 mg/kg), scopolamine (0.0056-1.0 mg/kg), nicotine (0.01-0.18 mg/kg), and guanfacine (0.0001-0.01 mg/kg) were dissolved in saline (0.9% sodium chloride + water). Donepezil was administered orally (p.o.) at a dose volume of 2.5 ml/kg, while all other drugs were administered through the intramuscular (i.m.) route with an injection volume of 0.1 ml/kg. Pre-treatment times were 4.5 hours for donepezil, 1 hour for scopolamine, 0.5 hours for nicotine and 2 hours for guanfacine. All drug doses were calculated from the free-base.

### Statistical Analysis

Two dependent measures were analyzed. “Hits” (i.e., accuracy) were defined as correct responses to targets, and the raw data were transformed into percent (percent hits = (number of hits/number of target trials) x 100). Reaction time was defined as the amount of time that elapsed between the appearance of the stimulus on the screen and the monkey’s touch response to the stimulus. An increase in accuracy (hits) indicated enhanced attention. Each dependent measure was analyzed with a one-way repeated measure ANOVA, and when necessary, the non-parametric equivalent (one-way repeated measure ANOVA on ranked data) was used. When a significant main effect was identified in the ANOVA, post hoc analysis was conducted by Fisher (non-parametric data) or Dunnett’s tests (parametric data), to determine at which particular dose(s) the effects were significant. Each variable was analyzed separately, and significance was defined as p < 0.05.

## RESULTS

### Scopolamine Induced Impairments on the CPT

The muscarinic acetylcholine receptor antagonist, scopolamine, was administered at a dose range of 0.0056-0.03 mg/kg to induce impairments on the CPT. There was a main effect of treatment with scopolamine for hits (F(4,23) = 18.19, p < 0.001; Figure 1a) and reaction times (F(4,22) = 16.37, p < 0.001; Figure 1b). For both dependent measures, post hoc analyses revealed that the 0.01, 0.018, and 0.03 mg/kg doses of scopolamine were significantly different from vehicle administration (all p < 0.05), indicating that scopolamine dose-dependently impaired CPT performance.

**Figure 1.**
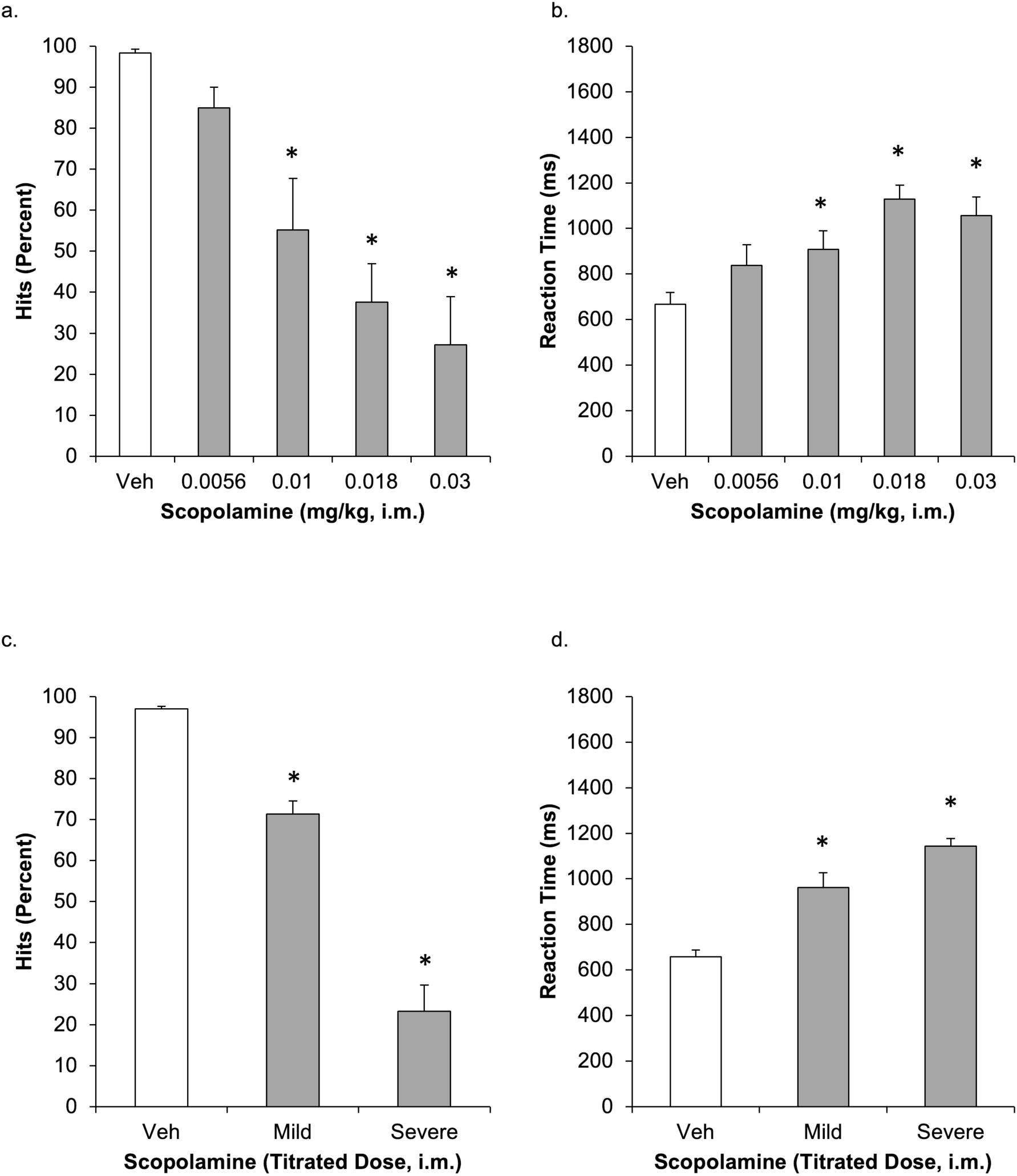
Acute scopolamine administration produced dose-dependent decreases in accuracy (% Hits) (a) and increases in reaction time to targets in NHPs performing the CPT (b). (c) and (d) depict the dose of scopolamine that impairs CPTS accuracy and reaction time, respectively, without impairing the ability of the animal to perform the task. Data are presented as group means ± SEM. *p < 0.05 vs vehicle.

To examine the effects of compounds with cognitive-enhancing properties, optimal scopolamine doses were selected for each monkey to produce a mildly impairing dose (71.4 ± 3.2% accuracy) and a severely impairing dose (23.3 ± 6.4% accuracy). Statistical analyses revealed a main effect of treatment for hits (F(2,17) = 73.38, p < 0.001; Figure 1c) and reaction times (F(2,17) = 64.35, p < 0.001; Figure 1d). Post hoc analyses indicated that both scopolamine doses significantly impaired CPT performance compared to vehicle (all p < 0.05).

### Guanfacine Reversal of Scopolamine-Induced Impairments

The selective α2A adrenergic agonist, guanfacine, was administered at a dose range of 0.0001-0.01 mg/kg following mildly or severely impairing doses of scopolamine to assess reversal of attentional impairments. Under conditions of mild impairment, guanfacine produced a significant main effect of treatment on hits (F(4,24) = 5.81, p = 0.002; Figure 2a) and reaction times (F(4,24) = 8.06, p < 0.001; Figure 2b). Post hoc analyses indicated that the mildly impairing dose of scopolamine significantly decreased hits compared to vehicle (p < 0.05), and the 0.0018, 0.003, and 0.0056 mg/kg doses of guanfacine significantly improved performance relative to scopolamine alone (all p < 0.05). Although scopolamine significantly increased reaction times compared to vehicle (p < 0.05), guanfacine did not significantly reverse this effect (p > 0.05).

**Figure 2.**
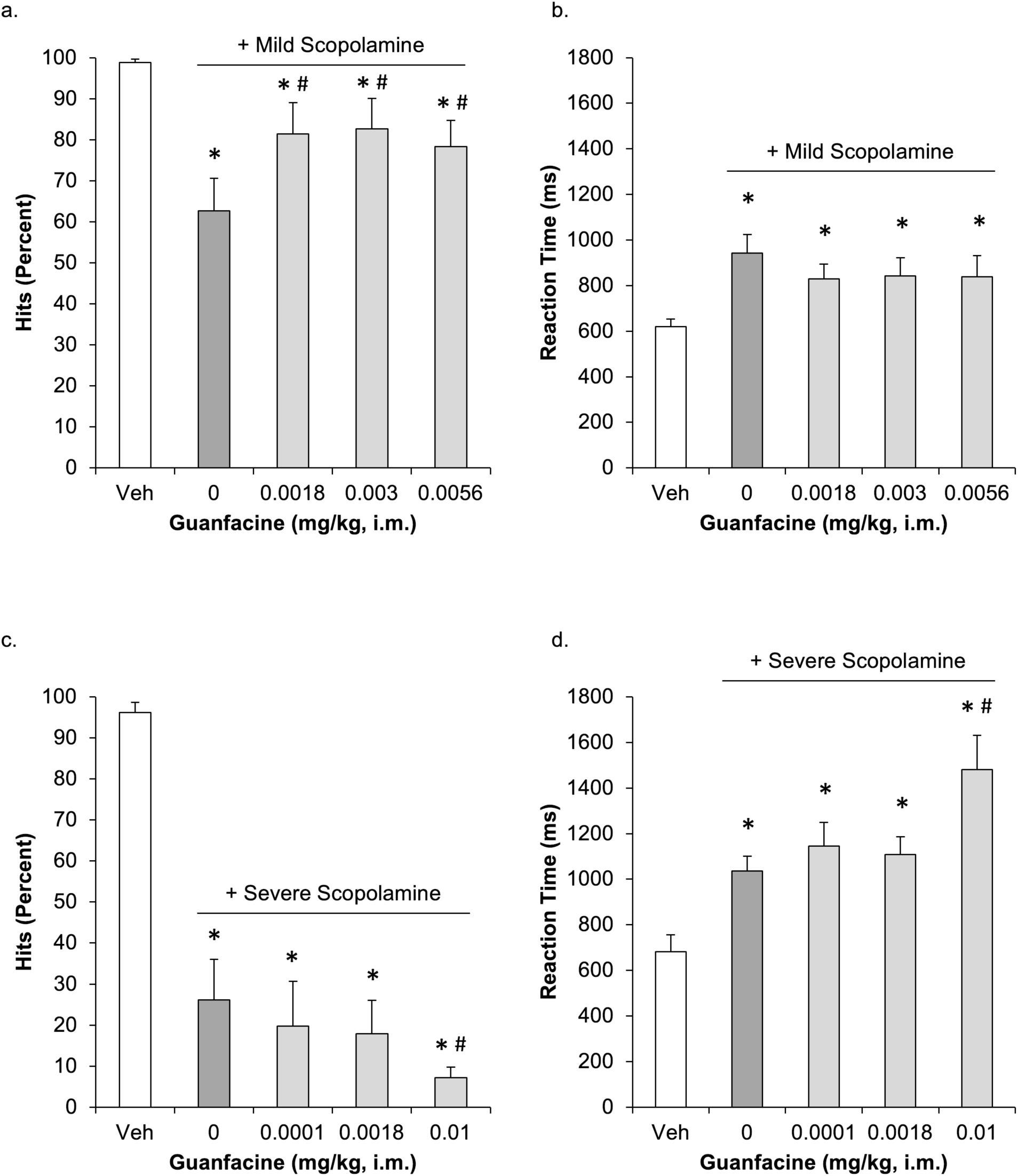
Acute guanfacine administration attenuated mild scopolamine-induced impairments in accuracy (% Hits) (a), but not severe scopolamine-induced impairments (c). Guanfacine did not attenuate either mild (b) or severe (d) scopolamine-induced slowing in reaction time. Data are presented as group means ± SEM. * p < 0.05 vs vehicle. ^#^p < 0.05 vs scopolamine alone.

Under conditions of severe impairment, guanfacine produced a significant main effect of treatment on hits (F(4,22) = 77.02, p < 0.001; Figure 2c) and reaction times (F(4,21) = 8.31, p < 0.001; Figure 2d(. Post hoc analyses confirmed that scopolamine significantly impaired performance (all p < 0.05). Notably, the 0.01 mg/kg dose of guanfacine further impaired performance on both measures compared to scopolamine alone (all p < 0.05).

### Donepezil Reversal of Scopolamine-Induced Impairments

We next tested whether the acetylcholinesterase inhibitor, donepezil, could reverse the scopolamine-induced CPT impairments. Donepezil was administered at doses of 0.3-3.0 mg/kg following mildly or severely impairing doses of scopolamine. Under mild impairment, donepezil produced a significant main effect on hits (Chi-square = 12.85, df = 4, p = 0.012; Figure 3a) and reaction times (F(4,24) = 8.80, p < 0.001; Figure 3b). Post hoc analyses indicated that scopolamine significantly impaired performance compared to vehicle (all p < 0.05). However, donepezil did not significantly reverse these impairments, with the 1.8 mg/kg dose not significantly different from either vehicle or scopolamine (all p > 0.05).

**Figure 3.**
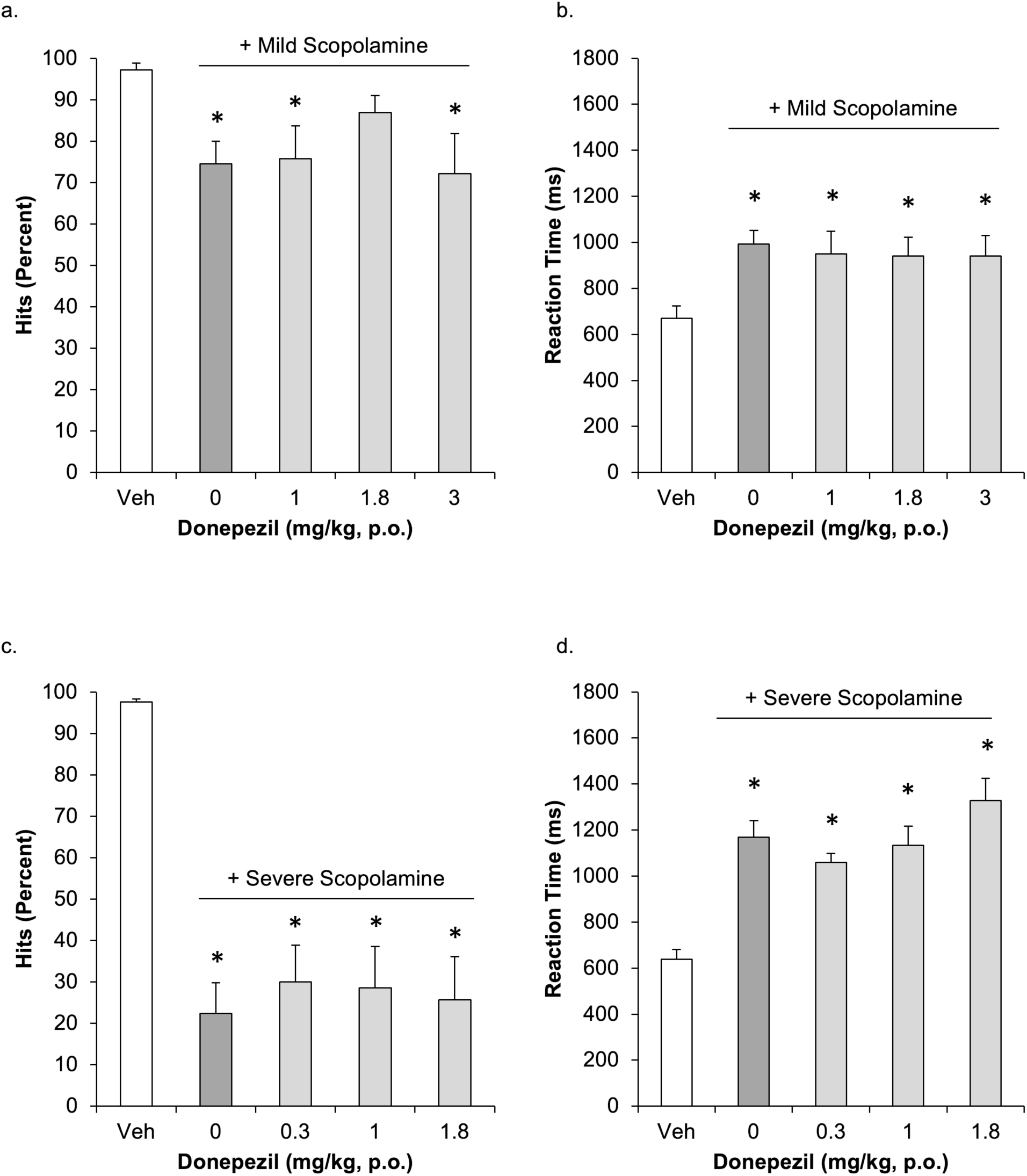
Acute donepezil administration did not attenuate mild (a) or severe (c) scopolamine- induced impairments in accuracy (%Hits) or reaction time (b and d). Data are presented as group means ± SEM. * p < 0.05 vs vehicle.

Under severe impairment, donepezil produced a significant main effect of treatment on hits (F(4,26) = 43.25, p < 0.001; Figure 3c) and reaction times (F(4,21) = 23.72, p < 0.001; Figure 3d). Post hoc analyses indicated significant impairment following scopolamine (all p < 0.05), but none of the donepezil doses significantly improved hits or reaction times (all p >0.05).

### Nicotine Reversal of Scopolamine-Induced Impairments

Finally, we administered the nicotinic acetylcholine receptor agonist, nicotine, at doses of 0.01-0.18 mg/kg following mildly or severely impairing doses of scopolamine. Under mild impairment, there was a significant main effect of treatment on hits (F(4,23) = 6.53, p = 0.001; Figure 4a) and reaction times (F(4,23) = 9.27, p < 0.001; Figure 4b). Post hoc analyses indicated that scopolamine significantly reduced hits compared to vehicle (p < 0.05), and the 0.03 mg/kg dose of nicotine significantly improved performance compared to scopolamine alone (p < 0.05). Nicotine did not reverse the scopolamine-induced slowing of reaction times (p > 0.05).

**Figure 4.**
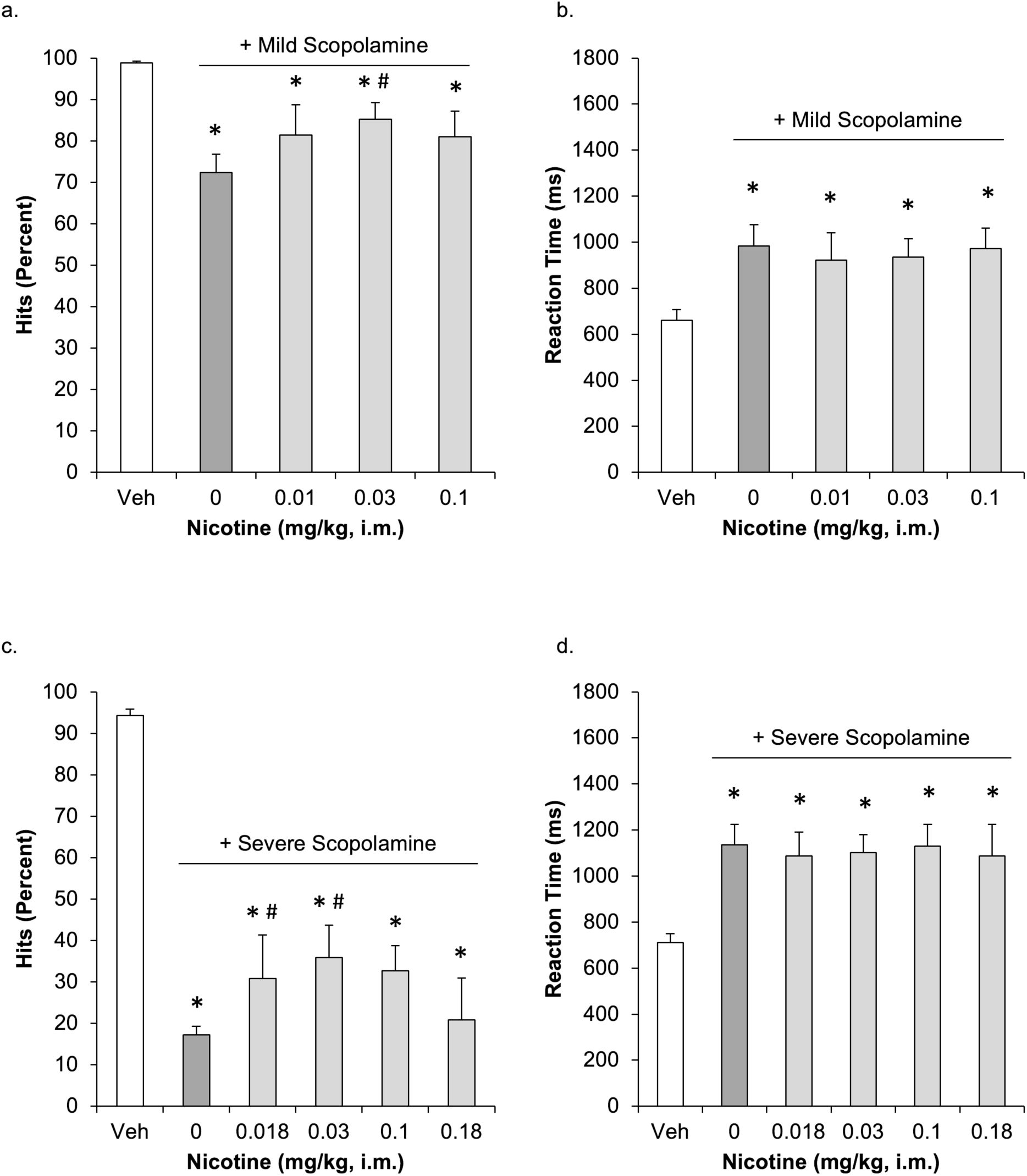
Acute nicotine administration attenuated both mild (a) and severe (c) scopolamine- induced impairments in accuracy (% Hits), but had no significant effect on reaction time under either impairing condition (b and d). Data are presented as group means ± SEM. * p < 0.05 vs vehicle.

Under severe impairment, nicotine produced a significant main effect of treatment on hits (F(5,25) = 35.49, p < 0.001; Figure 4c) and reaction times (F(5,25) = 3.68, p = 0.012; Figure 4d). Post hoc analyses indicated that scopolamine significantly decreased hits and increased reaction times (all p < 0.05). The 0.018 and 0.03 mg/kg doses of nicotine significantly increased hits compared to scopolamine alone (all p < 0.05), but had no significant effect on reaction times (p > 0.05).

## DISCUSSION

This study establishes a robust, translational model of sustained attention in NHPs using the CPT to investigate pharmacological reversal of attentional deficits. By inducing dose-dependent impairments with the muscarinic antagonist scopolamine, we successfully modeled both mild and severe cholinergic dysfunction, a hallmark of aging and AD. Our findings demonstrate that nicotine restored attention performance across mild and severe impairment levels, guanfacine was effective only under mild impairment, and donepezil yielded inconsistent results. These differential response patterns offer critical insight into the receptor-specific mechanisms underlying sustained attention and suggest that nicotinic agonists may be particularly well-suited for treating attentional impairments in both prodromal and more advanced stages of disease. Moreover, this work highlights the utility of the NHP CPT paradigm as a sensitive platform for probing the neuropharmacology of attention and for advancing targeted therapeutics for cognitive decline in aging and AD.

### Cholinergic Degeneration and Early Attention Deficits in Alzheimer’s Disease

Although AD is most commonly associated with memory loss, emerging evidence points to early cholinergic system dysfunction as a critical feature of the disease [47–50]. Sustained attention, the ability to maintain focus over time, is among the earliest non-memory domains to decline in AD [16,51–56,6]. A key contributor to these early changes is the degeneration of cholinergic neurons in the basal forebrain, particularly the nucleus basalis of Meynert, which provides the primary source of cortical cholinergic input [57–61]. Neurofibrillary tangle accumulation in this region correlates strongly with dementia severity [57–61]. While the PFC, which supports top-down attentional control, is affected later in the disease, its function may be compromised early due to the loss of cholinergic input [50,62–64]. These findings suggest that attentional decline is not simply secondary to memory dysfunction but represents a distinct and early cognitive marker of AD. As such, attentional impairments may offer both a sensitive indicator for early detection and a meaningful target for therapeutic intervention.

### Scopolamine as a Translational Tool for Modeling Cholinergic Dysfunction in Alzheimer’s Disease

Given the early and progressive involvement of the cholinergic system AD and its close association attentional deficits, pharmacological models that impair cholinergic signaling provide a valuable tool for probing these mechanisms in controlled experimental settings. Scopolamine, a non-selective muscarinic acetylcholine receptor antagonist, induces transient cholinergic disruption and reliably produces attention and memory impairments across species [65–67].

Scopolamine binds with high affinity to M1 and M2 muscarinic receptor subtypes [68] and can block nicotinic receptors at higher doses [69], making it particularly relevant for modeling the cholinergic deficits observed in AD.

In this study, individualized scopolamine doses were used to produce mild or severe impairments on the CPT, simulating early and later stages of cholinergic dysfunction. These two levels of impairment enabled the assessment of cognitive enhancers across a spectrum of severity, mirroring disease progression. Our finding that scopolamine impairs CPT performance aligns with extensive prior work in rodents, NHPs, and humans [70–79]. Notably, the level of impairment observed with mild scopolamine matched the performance of aged NHPs on the same task [35], supporting scopolamine’s translational validity. By simulating different stages of attentional dysfunction, this model provides a powerful and scalable platform for evaluating pro-cognitive agents in a manner relevant to human disease.

### Targeting the Noradrenergic System: Selective Effects of Guanfacine

Norepinephrine plays a critical role in modulating attention, with both insufficient and excessive levels disrupting PFC function [80,81]. In healthy individuals, LC provides the primary source of cortical norepinephrine. In AD, however, degeneration of the LC leads to a loss of up to 70% of noradrenergic neurons, contributing to early attentional decline [27]. At the cellular level, norepinephrine binds to α2A adrenergic receptors on post-synaptic dendritic spines, reducing cAMP signaling, closing HCN and potassium channels, and enhancing neuronal firing in the PFC. These actions strengthen top-down regulation of attention and executive control [33]. Guanfacine, a selective α2A receptor agonist, mimics these effects and has been shown to improve attentional performance in both aged humans and non-human primates [31,82,35,83].

In our study, guanfacine significantly improved CPT accuracy following mild scopolamine-induced impairment, consistent with prior work [35]. This enhancement was selective to attentional performance, as reaction time was unaffected, suggesting a specific effect on top-down control rather than general arousal. However, under severe impairment, guanfacine further worsened performance. Similar findings have been reported with α2 agonists such as clonidine in AD patients, where excessive suppression of LC activity may reduce cortical norepinephrine below functional thresholds [84]. These results highlight a narrow therapeutic window for noradrenergic agents and underscore the importance of neuromodulatory balance in PFC function.

### Limited Efficacy of Donepezil in Reversing Scopolamine-Induced Attention Deficits

A substantial body of literature has demonstrated that AD leads to degeneration of cholinergic neurons, resulting in widespread reduction in ACh signaling [85,86]. This loss is particularly pronounced in the basal forebrain, including the nucleus basalis of Meynert (NBM), which provides dense cholinergic innervation to the PFC [64]. In response, many AD therapies have focused on acetylcholinesterase inhibitors, such as donepezil, which aim to enhance cholinergic tone by preventing enzymatic breakdown of ACh in the synaptic cleft [87]. By increasing ACh availability, these compounds are hypothesized to compensate for presynaptic neuron loss and support residual cholinergic transmission, thereby mitigating cognitive dysfunction.

Donepezil, an acetylcholinesterase inhibitor approved for the treatment of all stages of AD, has shown benefits in domains such as memory and global cognition [88–90]. However, in our study, donepezil failed to significantly improve sustained attention on the CPT under either mild or severe cholinergic impairment. While there was a non-significant trend toward improvement under mild impairment, the effect did not reach statistical significance. These findings differ from prior studies showing cognitive benefits of donepezil in both animal models and clinical populations [91,92], though much of that work has focused on spatial or working memory rather than sustained attention.

Mechanistically, the limited efficacy observed here may reflect a key constraint of cholinesterase inhibition: its reliance on the presence of functional muscarinic receptors. When receptors are pharmacologically blocked, as with scopolamine, or physically degraded in advanced AD, elevated ACh levels alone may be insufficient to restore downstream signaling. This pharmacodynamic ceiling may explain why donepezil is inconsistently effective in clinical trials and why its benefits are often modest, particularly for attentional processes, which depend on finely tuned cholinergic activity in PFC circuits. Our findings reinforce the notion that cholinesterase inhibitors may have limited utility for non-memory domains such as sustained attention, especially in the context of more advanced cholinergic system degeneration.

### Nicotinic Agonism Enhances Sustained Attention Across Impairment Severity

Given the limitations of cholinesterase inhibitors, nicotinic acetylcholine receptors (nAChRs) have emerged as promising alternative targets for cognitive enhancement. Nicotine, a non-selective nAChR agonist, enhances attentional performance across species [93–97]. In our study, nicotine significantly improved CPT accuracy under both mild and severe cholinergic disruption, outperforming both guanfacine and donepezil. Critically, nicotine improved hit rates without affecting reaction time, suggesting a specific effect on sustained attention rather than general arousal or motor facilitation. Direct receptor agonism may allow nAChR engagement even when endogenous ACh is limited, offering a potential advantage over cholinesterase inhibitors in later disease stages. These results support ongoing interest in nicotinic receptor modulators as cognitive therapeutics and underscore the need to characterize the receptor subtypes driving these effects.

### Summary

Although guanfacine, donepezil, and nicotine varied in their efficacy, none of the tested compounds reversed the scopolamine-induced slowing of reaction times. This consistent dissociation supports the conclusion that the observed drug effects were specific to sustained attentional control rather than general enhancements in arousal or motor speed. Furthermore, our use of full dose-response curves, rather than single best-dose comparisons, increased the precision and translational relevance of the model. This approach captured variability across subjects and more closely mirrors the heterogeneity observed in human clinical populations.

**Table 1.** Dependent measures for all studies and conditions.

| Study | Dose | Dependent Measure |  |  |  |  |  |  |  |  |  |
| --- | --- | --- | --- | --- | --- | --- | --- | --- | --- | --- | --- |
|  |  | Hits |  | Misses |  | Correct Rejections |  | False Alarms |  | Reaction Time |  |
|  |  | Mean (%) | SEM | Mean (%) | SEM | Mean (%) | SEM | Mean (%) | SEM | Mean (ms) | SEM |
| Scopolamine<br>Dose-Response<br>Curve | Vehicle | 98.33 | 1.00 | 1.67 | 1.00 | 92.86 | 3.04 | 7.14 | 3.04 | 668.09 | 50.50 |
|  | 0.0056 | 85.00 | 4.97 | 15.00 | 4.97 | 98.57 | 0.75 | 1.43 | 0.75 | 837.06 | 92.10 |
|  | 0.01 | 55.21 | 12.59 | 44.79 | 12.59 | 95.09 | 1.76 | 4.91 | 1.76 | 907.27 | 83.25 |
|  | 0.018 | 37.59 | 9.41 | 62.41 | 9.41 | 98.65 | 0.77 | 1.35 | 0.77 | 1129.75 | 59.73 |
|  | 0.03 | 27.14 | 11.73 | 72.86 | 11.73 | 98.06 | 0.81 | 1.94 | 0.81 | 1056.80 | 81.54 |
| Scopolamine Mild<br>and Severe | Vehicle | 97.01 | 0.64 | 2.99 | 0.64 | 94.21 | 2.56 | 5.79 | 2.56 | 657.83 | 29.61 |
|  | Mild | 71.36 | 3.20 | 28.64 | 3.20 | 95.72 | 1.65 | 4.28 | 1.65 | 962.00 | 64.55 |
|  | Severe | 23.25 | 6.43 | 76.75 | 6.43 | 97.72 | 1.15 | 2.28 | 1.15 | 1143.23 | 34.72 |
| Mild Scopolamine<br>+<br>Guanfacine | Vehicle | 98.81 | 0.79 | 1.19 | 0.79 | 93.37 | 3.29 | 6.63 | 3.29 | 619.70 | 33.59 |
|  | Mild + 0 | 62.62 | 7.98 | 37.38 | 7.98 | 94.90 | 2.04 | 5.10 | 2.04 | 942.64 | 81.43 |
|  | Mild + 0.0018 | 81.43 | 7.62 | 18.57 | 7.62 | 96.94 | 1.27 | 3.06 | 1.27 | 828.58 | 65.76 |
|  | Mild + 0.003 | 82.62 | 7.40 | 17.38 | 7.40 | 98.16 | 0.79 | 1.84 | 0.79 | 842.59 | 79.07 |
|  | Mild + 0.0056 | 78.33 | 6.36 | 21.67 | 6.36 | 97.96 | 1.08 | 2.04 | 1.08 | 837.74 | 92.97 |
| Severe<br>Scopolamine<br>+<br>Guanfacine | Vehicle | 96.19 | 2.43 | 3.81 | 2.43 | 97.04 | 0.90 | 2.96 | 0.90 | 682.45 | 73.49 |
|  | Severe + 0 | 26.19 | 9.87 | 73.81 | 9.87 | 98.47 | 0.94 | 1.53 | 0.94 | 1035.40 | 65.89 |
|  | Severe + 0.0001 | 19.72 | 10.92 | 80.28 | 10.92 | 98.93 | 0.66 | 1.07 | 0.66 | 1144.69 | 105.47 |
|  | Severe + 0.0018 | 17.86 | 8.15 | 82.14 | 8.15 | 96.73 | 2.91 | 3.27 | 2.91 | 1108.57 | 77.77 |
|  | Severe + 0.01 | 7.14 | 2.64 | 92.86 | 2.64 | 99.49 | 0.34 | 0.51 | 0.34 | 1481.44 | 150.29 |
| Mild Scopolamine<br>+<br>Donepezil | Vehicle | 97.14 | 1.66 | 2.86 | 1.66 | 97.04 | 1.37 | 2.96 | 1.37 | 669.73 | 53.92 |
|  | Mild + 0 | 73.89 | 6.36 | 26.11 | 6.36 | 99.52 | 0.24 | 0.48 | 0.24 | 985.58 | 69.92 |
|  | Mild + 1 | 72.50 | 8.63 | 27.50 | 8.63 | 98.33 | 0.73 | 1.67 | 0.73 | 952.81 | 116.85 |
|  | Mild + 1.8 | 86.90 | 4.06 | 13.10 | 4.06 | 98.37 | 0.53 | 1.63 | 0.53 | 941.36 | 80.02 |
|  | Mild + 3 | 72.14 | 9.67 | 27.86 | 9.67 | 99.18 | 0.70 | 0.82 | 0.70 | 941.22 | 87.74 |
| Severe<br>Scopolamine<br>+<br>Donepezil | Vehicle | 97.59 | 0.69 | 2.41 | 0.69 | 98.17 | 0.73 | 1.83 | 0.73 | 638.52 | 42.82 |
|  | Severe + 0 | 22.41 | 7.39 | 77.59 | 7.39 | 96.67 | 1.28 | 3.33 | 1.28 | 1169.23 | 71.69 |
|  | Severe + 0.3 | 30.00 | 8.84 | 70.00 | 8.84 | 96.70 | 0.94 | 3.30 | 0.94 | 1059.82 | 39.20 |
|  | Severe + 1 | 28.54 | 9.98 | 71.46 | 9.98 | 98.04 | 1.12 | 1.96 | 1.12 | 1133.25 | 83.67 |
|  | Severe + 1.8 | 25.67 | 10.44 | 74.33 | 10.44 | 97.14 | 1.48 | 2.86 | 1.48 | 1327.20 | 97.59 |
| Mild Scopolamine<br>+<br>Nicotine | Vehicle | 98.81 | 0.48 | 1.19 | 0.48 | 94.80 | 3.33 | 5.20 | 3.33 | 661.46 | 45.31 |
|  | Mild + 0 | 72.38 | 4.35 | 27.62 | 4.35 | 98.27 | 0.73 | 1.73 | 0.73 | 983.06* | 93.12 |
|  | Mild + 0.01 | 81.39 | 7.36 | 18.61 | 7.36 | 97.14 | 1.74 | 2.86 | 1.74 | 922.56* | 118.12 |
|  | Mild + 0.03 | 85.24 | 4.01 | 14.76 | 4.01 | 98.06 | 1.71 | 1.94 | 1.71 | 934.31* | 80.88 |
|  | Mild + 0.1 | 84.05 | 5.57 | 15.95 | 5.57 | 98.78 | 0.81 | 1.22 | 0.81 | 927.47* | 95.05 |
| Severe<br>Scopolamine<br>+<br>Nicotine | Vehicle | 94.31 | 1.53 | 5.69 | 1.53 | 89.64 | 7.83 | 10.36 | 7.83 | 710.33 | 39.24 |
|  | Severe + 0 | 17.22 | 2.09 | 82.78 | 2.09 | 97.98 | 0.81 | 2.02 | 0.81 | 1135.90* | 89.02 |
|  | Severe + 0.018 | 30.83 | 10.46 | 69.17 | 10.46 | 93.81 | 4.91 | 6.19 | 4.91 | 1087.02* | 103.78 |
|  | Severe + 0.03 | 35.83 | 7.85 | 64.17 | 7.85 | 94.88 | 2.54 | 5.12 | 2.54 | 1101.85* | 77.17 |
|  | Severe + 0.1 | 32.64 | 6.09 | 67.36 | 6.09 | 93.81 | 3.17 | 6.19 | 3.17 | 1129.83* | 95.11 |
|  | Severe + 0.18 | 20.83 | 10.13 | 79.17 | 10.13 | 94.05 | 3.56 | 5.95 | 3.56 | 1087.11* | 137.18 |

\* p < 0.05 vs vehicle

In summary, we have demonstrated that the CPT is readily trained in NHPs and that pharmacologically induced impairments in performance parallel those observed in aged NHPs and are mechanistically relevant to disease processes. Additionally, we validated the model for testing cognitive-enhancing compounds across distinct neuromodulatory systems. Finally, we showed that dose-response curve analysis is both feasible and informative in this paradigm.

These findings identify nicotinic receptor activation as a particularly promising avenue for rescuing attentional deficits and establish the NHP CPT model as a scalable, translatable platform for evaluating pro-cognitive therapeutics. Future studies may build on this framework by dissecting the contributions of nicotinic receptor subtypes and expanding therapeutic screening to include compounds developed for related disorders such as schizophrenia and ADHD.

## Data Availability Statement

The datasets generated during the current study are available from the corresponding author on reasonable request.

### Acknowledgments

This work was supported by F. Hoffmann-La Roche.

## Author Contributions

TMB, JGW, and CGB contributed to the conception and design of the work, CGB and JSC collected the data, CRV, SRD, and CGB analyzed and interpreted the data and drafted the manuscript, all authors approved final version for publication.

## REFERENCES

1. Simon AJ, Gallen CL, Ziegler DA, Mishra J, Marco EJ, Anguera JA, et al. Quantifying attention span across the lifespan. Front Cogn. 2023;2.

2. Gazzaley A, Cooney JW, Rissman J, D’Esposito M. Top-down suppression deficit underlies working memory impairment in normal aging. Nat Neurosci. 2005;8:1298–1300.

3. Huntley JD, Hampshire A, Bor D, Owen AM, Howard RJ. The importance of sustained attention in early Alzheimer’s disease. Int J Geriatr Psychiatry. 2017;32:860–867.

4. Vanderlip CR, Stark CEL, Initiative for the ADN. Digital cognitive assessments as low-burden markers for predicting future cognitive decline and tau accumulation across the Alzheimer’s spectrum. Alzheimers Dement. 2024;20:6881–6895.

5. Malhotra PA. Impairments of attention in Alzheimer’s disease. Curr Opin Psychol. 2019;29:41–48.

6. Berardi AM, Parasuraman,Raja, and Haxby JV. Sustained Attention in Mild Alzheimer’s Disease. Dev Neuropsychol. 2005;28:507–537.

7. Ball KK, Ross LA, Roth DL, Edwards JD. Speed of processing training in the ACTIVE study: how much is needed and who benefits? J Aging Health. 2013;25:65S–84S.

8. Paneri S, Gregoriou GG. Top-Down Control of Visual Attention by the Prefrontal Cortex. Functional Specialization and Long-Range Interactions. Front Neurosci. 2017;11.

9. Rossi AF, Pessoa L, Desimone R, Ungerleider LG. The prefrontal cortex and the executive control of attention. Exp Brain Res Exp Hirnforsch Exp Cerebrale. 2009;192:489–497.

10. Cohen RM, Semple WE, Gross M, Nordahl TE, Holcomb HH, Dowling MS, et al. The effect of neuroleptics on dysfunction in a prefrontal substrate of sustained attention in schizophrenia. Life Sci. 1988;43:1141–1150.

11. Pardo JV, Fox PT, Raichle ME. Localization of a human system for sustained attention by positron emission tomography. Nature. 1991;349:61–64.

12. Posner MI, Petersen SE. The attention system of the human brain. Annu Rev Neurosci. 1990;13:25–42.

13. Coull JT, Frith CD, Frackowiak RS, Grasby PM. A fronto-parietal network for rapid visual information processing: a PET study of sustained attention and working memory. Neuropsychologia. 1996;34:1085–1095.

14. Lewin JS, Friedman L, Wu D, Miller DA, Thompson LA, Klein SK, et al. Cortical Localization of Human Sustained Attention: Detection with Functional MR Using a Visual Vigilance Paradigm. J Comput Assist Tomogr. 1996;20:695.

15. Davis RC, Maloney MT, Minamide LS, Flynn KC, Stonebraker MA, Bamburg JR. Mapping cofilin-actin rods in stressed hippocampal slices and the role of cdc42 in amyloid-beta-induced rods. J Alzheimers Dis JAD. 2009;18:35–50.

16. Perry RJ, Hodges JR. Attention and executive deficits in Alzheimer’s disease. A critical review. Brain J Neurol. 1999;122 (Pt 3):383–404.

17. Raz N, Lindenberger U, Rodrigue KM, Kennedy KM, Head D, Williamson A, et al. Regional Brain Changes in Aging Healthy Adults: General Trends, Individual Differences and Modifiers. Cereb Cortex. 2005;15:1676–1689.

18. Mesulam M. The cholinergic lesion of Alzheimer’s disease: pivotal factor or side show? Learn Mem Cold Spring Harb N. 2004;11:43–49.

19. Mesulam MM, Geula C. Acetylcholinesterase-rich pyramidal neurons in the human neocortex and hippocampus: absence at birth, development during the life span, and dissolution in Alzheimer’s disease. Ann Neurol. 1988;24:765–773.

20. Muir JL, Everitt BJ, Robbins TW. Reversal of visual attentional dysfunction following lesions of the cholinergic basal forebrain by physostigmine and nicotine but not by the 5-HT3 receptor antagonist, ondansetron. Psychopharmacology (Berl). 1995;118:82–92.

21. Voytko ML, Olton DS, Richardson RT, Gorman LK, Tobin JR, Price DL. Basal forebrain lesions in monkeys disrupt attention but not learning and memory. J Neurosci Off J Soc Neurosci. 1994;14:167–186.

22. Robbins TW, Granon S, Muir JL, Durantou F, Harrison A, Everitt BJ. Neural systems underlying arousal and attention. Implications for drug abuse. Ann N Y Acad Sci. 1998;846:222– 237.

23. Passetti F, Dalley JW, O’Connell MT, Everitt BJ, Robbins TW. Increased acetylcholine release in the rat medial prefrontal cortex during performance of a visual attentional task. Eur J Neurosci. 2000;12:3051–3058.

24. Dalley JW, McGaughy J, O’Connell MT, Cardinal RN, Levita L, Robbins TW. Distinct changes in cortical acetylcholine and noradrenaline efflux during contingent and noncontingent performance of a visual attentional task. J Neurosci Off J Soc Neurosci. 2001;21:4908–4914.

25. Schliebs R, Arendt T. The cholinergic system in aging and neuronal degeneration. Behav Brain Res. 2011;221:555–563.

26. Sahakian BJ, Owen AM, Morant NJ, Eagger SA, Boddington S, Crayton L, et al. Further analysis of the cognitive effects of tetrahydroaminoacridine (THA) in Alzheimer’s disease: assessment of attentional and mnemonic function using CANTAB. Psychopharmacology (Berl). 1993;110:395–401.

27. Berridge CW, Waterhouse BD. The locus coeruleus-noradrenergic system: modulation of behavioral state and state-dependent cognitive processes. Brain Res Brain Res Rev. 2003;42:33–84.

28. Arnsten AF. Catecholamine mechanisms in age-related cognitive decline. Neurobiol Aging. 1993;14:639–641.

29. Brozoski TJ, Brown RM, Rosvold HE, Goldman PS. Cognitive deficit caused by regional depletion of dopamine in prefrontal cortex of rhesus monkey. Science. 1979;205:929–932.

30. Carli M, Robbins TW, Evenden JL, Everitt BJ. Effects of lesions to ascending noradrenergic neurones on performance of a 5-choice serial reaction task in rats; implications for theories of dorsal noradrenergic bundle function based on selective attention and arousal. Behav Brain Res. 1983;9:361–380.

31. Arnsten AF, Contant TA. Alpha-2 adrenergic agonists decrease distractibility in aged monkeys performing the delayed response task. Psychopharmacology (Berl). 1992;108:159– 169.

32. O’Neill J, Fitten LJ, Siembieda DW, Ortiz F, Halgren E. Effects of guanfacine on three forms of distraction in the aging macaque. Life Sci. 2000;67:877–885.

33. Wang M, Ramos BP, Paspalas CD, Shu Y, Simen A, Duque A, et al. Alpha2A- adrenoceptors strengthen working memory networks by inhibiting cAMP-HCN channel signaling in prefrontal cortex. Cell. 2007;129:397–410.

34. Grudzien A, Shaw P, Weintraub S, Bigio E, Mash DC, Mesulam MM. Locus coeruleus neurofibrillary degeneration in aging, mild cognitive impairment and early Alzheimer’s disease. Neurobiol Aging. 2007;28:327–335.

35. Decamp E, Clark K, Schneider JS. Effects of the Alpha-2 Adrenoceptor Agonist Guanfacine on Attention and Working Memory in Aged Non-Human Primates. Eur J Neurosci. 2011;34:1018–1022.

36. Preuss TM. Do rats have prefrontal cortex? The rose-woolsey-akert program reconsidered. J Cogn Neurosci. 1995;7:1–24.

37. Wise SP. Forward frontal fields: phylogeny and fundamental function. Trends Neurosci. 2008;31:599–608.

38. Robbins TW, Arnsten AFT. The neuropsychopharmacology of fronto-executive function: monoaminergic modulation. Annu Rev Neurosci. 2009;32:267–287.

39. Porrino LJ, Goldman-Rakic PS. Brainstem innervation of prefrontal and anterior cingulate cortex in the rhesus monkey revealed by retrograde transport of HRP. J Comp Neurol. 1982;205:63–76.

40. Petrides M. Lateral prefrontal cortex: architectonic and functional organization. Philos Trans R Soc Lond B Biol Sci. 2005;360:781–795.

41. Beck LH, Bransome ED, Mirsky AF, Rosvold HE, Sarason I. A continuous performance test of brain damage. J Consult Psychol. 1956;20:343–350.

42. de Frias CM, Bunce D, Wahlin A, Adolfsson R, Sleegers K, Cruts M, et al. Cholesterol and triglycerides moderate the effect of apolipoprotein E on memory functioning in older adults. J Gerontol B Psychol Sci Soc Sci. 2007;62:P112–118.

43. Riccio CA, Reynolds CR, Lowe P, Moore JJ. The continuous performance test: a window on the neural substrates for attention? Arch Clin Neuropsychol. 2002;17:235–272.

44. Goonawardena AV, Heiss J, Glavis-Bloom C, Trube G, Borroni E, Alberati D, et al. Alterations in High-Frequency Neuronal Oscillations in a Cynomolgus Macaque Test of Sustained Attention Following NMDA Receptor Antagonism. Neuropsychopharmacology. 2016;41:1319–1328.

45. Zeamer A, Decamp E, Clark K, Schneider JS. Attention, executive functioning and memory in normal aged rhesus monkeys. Behav Brain Res. 2011;219:23–30.

46. Golub MS, Hogrefe CE, Germann SL, Tran TT, Beard JL, Crinella FM, et al. Neurobehavioral evaluation of rhesus monkey infants fed cow’s milk formula, soy formula, or soy formula with added manganese. Neurotoxicol Teratol. 2005;27:615–627.

47. Ferreira-Vieira TH, Guimaraes IM, Silva FR, Ribeiro FM. Alzheimer’s Disease: Targeting the Cholinergic System. Curr Neuropharmacol. 2016;14:101–115.

48. Hampel H, Mesulam M-M, Cuello AC, Farlow MR, Giacobini E, Grossberg GT, et al. The cholinergic system in the pathophysiology and treatment of Alzheimer’s disease. Brain. 2018;141:1917–1933.

49. Richter N, Beckers N, Onur OA, Dietlein M, Tittgemeyer M, Kracht L, et al. Effect of cholinergic treatment depends on cholinergic integrity in early Alzheimer’s disease. Brain. 2018;141:903–915.

50. Mufson EJ, Counts SE, Perez SE, Ginsberg SD. Cholinergic system during the progression of Alzheimer’s disease: therapeutic implications. Expert Rev Neurother. 2008;8:1703–1718.

51. Alexander DA. Attention dysfunction in senile dementia. Psychol Rep. 1973;32:229–230.

52. Sahakian B, Jones G, Levy R, Gray J, Warburton D. The effects of nicotine on attention, information processing, and short-term memory in patients with dementia of the Alzheimer type. Br J Psychiatry J Ment Sci. 1989;154:797–800.

53. Jones GM, Sahakian BJ, Levy R, Warburton DM, Gray JA. Effects of acute subcutaneous nicotine on attention, information processing and short-term memory in Alzheimer’s disease. Psychopharmacology (Berl). 1992;108:485–494.

54. Lines CR, Dawson C, Preston GC, Reich S, Foster C, Traub M. Memory and attention in patients with senile dementia of the Alzheimer type and in normal elderly subjects. J Clin Exp Neuropsychol. 1991;13:691–702.

55. Grady CL, Haxby,J. V., Horwitz,B., Sundaram,M., Berg,G., Schapiro,M., et al. Longitudinal study of the early neuropsychological and cerebral metabolic changes in dementia of the Alzheimer type. J Clin Exp Neuropsychol. 1988;10:576–596.

56. Sunderland T, Tariot PN, Cohen RM, Weingartner H, Mueller EA, Murphy DL. Anticholinergic sensitivity in patients with dementia of the Alzheimer type and age-matched controls. A dose-response study. Arch Gen Psychiatry. 1987;44:418–426.

57. Arnold SE, Hyman BT, Flory J, Damasio AR, Van Hoesen GW. The topographical and neuroanatomical distribution of neurofibrillary tangles and neuritic plaques in the cerebral cortex of patients with Alzheimer’s disease. Cereb Cortex N Y N 1991. 1991;1:103–116.

58. Vogt BA, Crino PB, Vogt LJ. Reorganization of cingulate cortex in Alzheimer’s disease: neuron loss, neuritic plaques, and muscarinic receptor binding. Cereb Cortex N Y N 1991. 1992;2:526–535.

59. Samuel W, Masliah E, Hill LR, Butters N, Terry R. Hippocampal connectivity and Alzheimer’s dementia: effects of synapse loss and tangle frequency in a two-component model. Neurology. 1994;44:2081–2088.

60. Liu AKL, Chang RC-C, Pearce RKB, Gentleman SM. Nucleus basalis of Meynert revisited: anatomy, history and differential involvement in Alzheimer’s and Parkinson’s disease. Acta Neuropathol (Berl). 2015;129:527–540.

61. Kuhn J, Hardenacke K, Lenartz D, Gruendler T, Ullsperger M, Bartsch C, et al. Deep brain stimulation of the nucleus basalis of Meynert in Alzheimer’s dementia. Mol Psychiatry. 2015;20:353–360.

62. Reed BR, Jagust WJ, Seab JP. Mental status as a predictor of daily function in progressive dementia. The Gerontologist. 1989;29:804–807.

63. Pepeu G, Grazia Giovannini M. The fate of the brain cholinergic neurons in neurodegenerative diseases. Brain Res. 2017;1670:173–184.

64. Auld DS, Kornecook TJ, Bastianetto S, Quirion R. Alzheimer’s disease and the basal forebrain cholinergic system: relations to beta-amyloid peptides, cognition, and treatment strategies. Prog Neurobiol. 2002;68:209–245.

65. Buccafusco JJ, Terry AV, Webster SJ, Martin D, Hohnadel EJ, Bouchard KA, et al. The scopolamine-reversal paradigm in rats and monkeys: the importance of computer-assisted operant-conditioning memory tasks for screening drug candidates. Psychopharmacology (Berl). 2008;199:481–494.

66. Broks P, Preston GC, Traub M, Poppleton P, Ward C, Stahl SM. Modelling dementia: effects of scopolamine on memory and attention. Neuropsychologia. 1988;26:685–700.

67. Lenz RA, Baker JD, Locke C, Rueter LE, Mohler EG, Wesnes K, et al. The scopolamine model as a pharmacodynamic marker in early drug development. Psychopharmacology (Berl). 2012;220:97–107.

68. Hulme EC, Birdsall NJ, Burgen AS, Mehta P. The binding of antagonists to brain muscarinic receptors. Mol Pharmacol. 1978;14:737–750.

69. Schmeller T, Sporer F, Sauerwein M, Wink M. Binding of tropane alkaloids to nicotinic and muscarinic acetylcholine receptors. Pharm. 1995;50:493–495.

70. Drachman DA, Leavitt J. Human memory and the cholinergic system. A relationship to aging? Arch Neurol. 1974;30:113–121.

71. Bartus RT, Johnson HR. Short-term memory in the rhesus monkey: disruption from the anti-cholinergic scopolamine. Pharmacol Biochem Behav. 1976;5:39–46.

72. Dunnett SB. Comparative effects of cholinergic drugs and lesions of nucleus basalis or fimbria-fornix on delayed matching in rats. Psychopharmacology (Berl). 1985;87:357–363.

73. Bushnell PJ. Modelling working and reference memory in rats: Effects of scopolamine on delayed matching-to-position. Behav Pharmacol. 1990;1:419–427.

74. Robbins TW, Semple J, Kumar R, Truman MI, Shorter J, Ferraro A, et al. Effects of scopolamine on delayed-matching-to-sample and paired associates tests of visual memory and learning in human subjects: comparison with diazepam and implications for dementia. Psychopharmacology (Berl). 1997;134:95–106.

75. Taffe MA, Weed MR, Gold LH. Scopolamine alters rhesus monkey performance on a novel neuropsychological test battery. Cogn Brain Res. 1999;8:203–212.

76. Duka T, Ott H, Rohloff A, Voet B. The effects of a benzodiazepine receptor antagonist β- carboline ZK-93426 on scopolamine-induced impairment on attention, memory and psychomotor skills. Psychopharmacology (Berl). 1996;123:361–373.

77. Brazell C, Preston GC, Ward C, Lines CR, Traub M. The scopolamine model of dementia: chronic transdermal administration. J Psychopharmacol Oxf Engl. 1989;3:76–82.

78. Klinkenberg I, Blokland A. The validity of scopolamine as a pharmacological model for cognitive impairment: A review of animal behavioral studies. Neurosci Biobehav Rev. 2010;34:1307–1350.

79. Miravalles C, Cannon DM, Hallahan B. The effect of scopolamine on memory and attention: a systematic review and meta-analysis. Eur Psychiatry. 2025;68:e50.

80. Arnsten AFT. Norepinephrine and cognitive disorders. Brain Norepinephrine Neurobiol. Ther., New York, NY, US: Cambridge University Press; 2007. p. 408–435.

81. Gamo NJ, Wang M, Arnsten AFT. Methylphenidate and atomoxetine enhance prefrontal function through α2-adrenergic and dopamine D1 receptors. J Am Acad Child Adolesc Psychiatry. 2010;49:1011–1023.

82. Ramos BP, Stark D, Verduzco L, van Dyck CH, Arnsten AFT. Alpha2A-adrenoceptor stimulation improves prefrontal cortical regulation of behavior through inhibition of cAMP signaling in aging animals. Learn Mem Cold Spring Harb N. 2006;13:770–776.

83. Arnsten AFT, Jin LE. Guanfacine for the treatment of cognitive disorders: a century of discoveries at Yale. Yale J Biol Med. 2012;85:45–58.

84. Riekkinen M, Laakso MP, Jäkälä P. Clonidine impairs sustained attention and memory in Alzheimer’s disease. Neuroscience. 1999;92:975–982.

85. Chen Z-R, Huang J-B, Yang S-L, Hong F-F. Role of Cholinergic Signaling in Alzheimer’s Disease. Molecules. 2022;27:1816.

86. Hampel H, Mesulam M-M, Cuello AC, Farlow MR, Giacobini E, Grossberg GT, et al. The cholinergic system in the pathophysiology and treatment of Alzheimer’s disease. Brain. 2018;141:1917–1933.

87. Rinne JO, Kaasinen V, Järvenpää T, Någren K, Roivainen A, Yu M, et al. Brain acetylcholinesterase activity in mild cognitive impairment and early Alzheimer’s disease. J Neurol Neurosurg Psychiatry. 2003;74:113–115.

88. Kumar A, Gupta V, Sharma S. Donepezil. StatPearls, Treasure Island (FL): StatPearls Publishing; 2025.

89. Birks JS, Harvey RJ. Donepezil for dementia due to Alzheimer’s disease. Cochrane Database Syst Rev. 2018;2018:CD001190.

90. Seltzer B, Zolnouni P, Nunez M, Goldman R, Kumar D, Ieni J, et al. Efficacy of Donepezil in Early-Stage Alzheimer Disease: A Randomized Placebo-Controlled Trial. Arch Neurol. 2004;61:1852–1856.

91. Buccafusco JJ, Terry AV. Donepezil-induced improvement in delayed matching accuracy by young and old rhesus monkeys. J Mol Neurosci MN. 2004;24:85–91.

92. Birks J, Harvey RJ. Donepezil for dementia due to Alzheimer’s disease. Cochrane Database Syst Rev. 2006:CD001190.

93. Wallace TL, Porter RHP. Targeting the nicotinic alpha7 acetylcholine receptor to enhance cognition in disease. Biochem Pharmacol. 2011;82:891–903.

94. Ruiz NA, Thieu MK, Aly M. Cholinergic modulation of hippocampally mediated attention and perception. Behav Neurosci. 2021;135:51–70.

95. Gil SM, Metherate R. Enhanced Sensory–Cognitive Processing by Activation of Nicotinic Acetylcholine Receptors. Nicotine Tob Res. 2019;21:377–382.

96. Valentine G, Sofuoglu M. Cognitive Effects of Nicotine: Recent Progress. Curr Neuropharmacol. 2018;16:403–414.

97. Semenova S, Stolerman IP, Markou A. Chronic nicotine administration improves attention while nicotine withdrawal induces performance deficits in the 5-choice serial reaction time task in rats. Pharmacol Biochem Behav. 2007;87:360–368.

